# A highly sensitive and specific SYBR Green quantitative polymerase chain reaction (qPCR) method for rapid detection of scale drop disease virus in Asian sea bass, *Lates calcarifer*

**DOI:** 10.1101/849661

**Authors:** Sukhontip Sriisan, Chuenchit Boonchird, Siripong Thitamadee, Molruedee Sonthi, Ha Thanh Dong, Saengchan Senapin

## Abstract

Scale drop disease virus (SDDV) is a novel *Megalocytivirus* causing scale drop disease (SDD) in Asian sea bass in Southeast Asia. In order to support disease diagnosis and surveillance, the present study developed a highly sensitive and specific SYBR Green qPCR assay for rapid detection of SDDV. Specific primers targeting a 135-bp fragment of *ATPase* coding gene of the SDDV genome were newly designed and subsequent gradient PCR assays were conducted to investigate their optimal annealing temperature. The optimized qPCR assay could detect as low as 2 viral copies per reaction and showed no cross amplification with DNA extracted from 12 viruses and bacteria commonly found in aquatic animals. The SDDV *ATPase* qPCR method was subsequently validated with field samples (n= 86). The results revealed that all clinically sick fish (n=34) from 5 affected farms gave positive results. Interestingly, 30/52 samples of apparently healthy fish from 8 unaffected farms which previously tested negative for SDDV by semi-nested PCR assay were positive by the newly developed qPCR method. This suggested that qPCR method is highly sensitive and suitable for early screening of SDDV from clinically healthy fish and for disease confirmation of sick fish. Investigation of tissue tropism and viral load of SDDV revealed systemic viral infection with relatively high viral load (8 × 10^2^ to 6.8 × 10^4^ copies per 200 ng of DNA template) in all 9 tested organs including eyes, brain, fin, gills, kidney, liver, kidney, spleen, and muscle. The newly developed qPCR method in this study delivered an accurate and reliable method for rapid detection of SDDV that may facilitate active surveillance and prevent widespread of the virus.

**Highlights:**

- This study developed a SYBR Green qPCR assay for rapid detection of SDDV
- The developed qPCR assay is specific for SDDV with limit of detection of 2 viral copies per reaction
- The assay could detect the virus from subclinically infected fish with low viral load
- We recommend this qPCR assay for active surveillance and early screening of SDDV

## 1. Introduction

Scale drop syndrome (SDS) in Asian sea bass, *Lates calcarifer*, was first reported in Southeast Asia by Gibson-Kueh et al. (2012). The clinical symptoms of the diseased fish were characterized by darkened bodies, scale loss, tail and fin erosion, gills pallor, and sometimes exophthalmia (Gibson-Kueh et al., 2012). The cumulative mortality was estimated around 40-50% in natural disease outbreaks. Subsequently, the causative agent of SDS was identified as scale drop disease virus (SDDV), a novel member of the genus *Megalocytivirus* (de Groof et al., 2015). Based on transmission electron microscope, SDDV virions are icosahedral with diameter of approximately 140 nm, which is a common characteristic of viruses in the family *Iridoviridae*. SDDV is a double-stranded DNA virus with known incomplete genome of about 124 kb containing at least 129 open reading frames (de Groof et al., 2015).

The disease has been reported in a number of countries in Southeast Asia recently including Malaysia, Singapore, Indonesia and Thailand (de Groof et al., 2015; Senapin et al., 2019). However, presence of the virus in the region was linked to mortalities since 1992 with similar clinical signs but probably misdiagnosed as other pathogen infection(s) (de Groof et al., 2015). Rapid detection method is critically important for disease diagnosis and selection of SDDV-free fish for aquaculture as well as prevention of widespread of the virus. Up-to-date, several PCR methods have been developed including patented conventional PCR and probe-based qPCR protocols (Guelen et al., 2014; de Groof et al., 2015), publicly accessible semi-nested PCR and loop-mediated isothermal amplification (LAMP) assay (Charoenwai et al., 2019; Dangtip et al., 2019). This study aimed to develop a highly sensitive SYBR Green-based qPCR method for the specific detection of SDDV from not only clinically sick fish but also inapprently infected fish, in order to facilitates active surveillance program for Asian sea bass farming countries in Southeast Asia and elsewhere.

## 2. Materials and methods

### 2.1 DNA samples from Asian sea bass

DNA samples from kidney or liver of Asian sea bass (*Lates calcarifer*) with or without clinical signs of scale drop disease were obtained from our previous studies (Senapin et al., 2019; Charoenwai et al., 2019). Fish samples were collected from different farms during 2016-2018. Additional 3 samples with scale drop clinical signs were recently obtained in 2019 and included in the present study. Thus, a total number of 86 fish DNA samples were used in this study as listed in Table 1.

**Table 1.** Details of fish samples used in this study and the test results

| Year | Farm | Fish health status | Positive sample/Number of tested samples |  |
| --- | --- | --- | --- | --- |
|  |  |  | Semi-nested PCR* | qPCR |
| 2016 | Farm 1 | Sick fish with scale drop symptoms | 18/18 | 18/18<sup>#</sup> |
|  | Farm 2 | Unknown diseased fish | 6/6 | 6/6 |
| 2017 | Farm 3 | Sick fish with scale drop symptoms | 5/5 | 5/5 |
|  | Farm 4 | Clinically healthy fish | 0/12 | 8/12 |
|  | Farm 5 | Clinically healthy fish | 0/80 | 1/5 |
|  | Farm 6 | Clinically healthy fish | 0/10 | 0/5 |
|  | Farm 6 | Clinically healthy fish | 0/10 | 5/5 |
|  | Farm 8 | Clinically healthy fish | 0/10 | 1/5<sup>#</sup> |
|  | Farm 9 | Clinically healthy fish | 0/10 | 5/5<sup>#</sup> |
|  | Farm 10 | Clinically healthy fish | 0/5 | 5/5<sup>#</sup> |
|  | Farm 11 | Clinically healthy fish | 0/10 | 2/5<sup>#</sup> |
|  | Farm 12 | Clinically healthy fish | 0/13 | 3/5 |
| 2018 | Farm 13 | Sick fish with scale drop symptoms | 2/2 | 2/2 |
| 2019 | Farm 14 | Sick fish with scale drop symptoms | Not done | 3/3 |
| **Total** |  |  | **31/191**<br>**(16.2%)** | **64/86**<br>**(74.4%)** |

\*, previous results from Charoenwai et al., 2019
<sup>#</sup>, one positive amplicon from indicated sample set was subjected to DNA sequence analysis

### 2.2 Primer design

qPCR primers for SDDV detection were designed based on an *ATPase* (adenosine triphosphatase) coding gene of the SDDV genome sequences in the GenBank database using Primer 3 online software (Untergasser et al., 2012; Koressaar et al., 2007). Primers qSDDV-AF: 5’-AAT GAC CGA AAT ACG ACC GAG AAC −3’ and qSDDV-AR: 5’-GCG GGG ATC AAA TGT CGT TTT G-3’ yielding an amplicon of 135 bp (Fig. S1) were synthesized by Bio Basic Inc. Specificity of the primers was preliminarily tested *in silico* using Primer-BLAST program (https://www.ncbi.nlm.nih.gov/tools/primer-blast/).

### 2.3 Gradient PCR assay

To determine the optimal temperature of the designed qPCR primers, gradient PCR reactions were performed using annealing temperature ranging from 55 to 65 °C. The reaction mixture of 20 μl contained 200 ng of DNA extracted from SDDV-infected fish, 1x PCRBIO buffer (containing 3 mM MgCl2, 1 mM dNTPs, and manufacturer’s enhancer and stabilizers), 0.25 unit of PCRBIO *Taq* polymerase (PCRBIOSYSTEMS, cat.no. PB10.11-05) and 150 μM of each primer. The cycling conditions consisted of denaturation at 95°C for 30 s followed by 30 cycles of 55-65 °C for 15 s and 72°C for 15 s (Biometra machine). Gradient PCR products were then analyzed by agarose gel electrophoresis.

### 2.4 DNA cloning and sequence analysis

Expected products of SDDV *ATPase* fragment amplification (135 bp) obtained from the above PCR assays were cloned into pGEM-T easy vector (Promega) and transformed into *E. coli* XL-1 Blue competent cells. Recombinant plasmid with the desired insert size was sent for DNA sequencing by Macrogen (South Korea). Plasmid DNA was used as positive control and used in qPCR sensitivity assay. Representative qPCR amplicons later found from positive test samples were also subjected to DNA cloning and sequencing in the same manner. Multiple sequence alignment of the obtained sequences was conducted using Clustal Omega (Sievers et al., 2011).

### 2.5 Optimization of SDDV qPCR conditions

The SDDV qPCR assay was performed using KAPA SYBR FAST Master Mix ABI Prism (KAPABIOSYSTEMS, cat. no. KM4603). A reaction of 20 μl contained 1x master mix, 150 or 200 nM of each forward and reverse primer, 2 μl of DNA template and molecular-grade water to adjust the final volume. The cycling conditions consisted of denaturation at 95°C for 3 min followed by 40 cycles of 95°C for 3 s and 60 or 63°C for 30 s (ABI 7500 instrument). At the end of the PCR amplification, melting curve analysis was performed.

### 2.6 Efficiency of amplification, detection sensitivity, and reproducibility

The optimized qPCR conditions obtained above were subjected to investigation of i) the efficiency of amplification (E) of the designed primer pair, ii) detection sensitivity, and iii) reproducibility test by intra-and inter-assays. These were done using 10-fold serial dilutions of the SDDV *ATPase* control plasmid. The concentration of plasmid used was ranged from 2 to 2×10^6^ copies per reaction. Additionally, 100 ng of DNA extracted from SDDV-free Asian sea bass was spiked in each qPCR reaction to mimic real detection assays. Standard curves were prepared by plotting the log of concentrations of plasmid versus threshold cycle (Ct) values. Amplification efficiency, E was then determined using the slope of the graph with the equation E = 10[^−1/slope^] (Pfaffl, 2001). With respect to detection sensitivity assay, detection limit of SDDV qPCR assay was determined from the minimum copy number that can could still be detected. For reproducibility test, the intra-and inter-assays were performed in 3 replicates.

### 2.7 Specificity test

Specificity test of the newly developed SDDV qPCR assay was performed against DNA samples prepared from 12 pathogens commonly found in aquatic animals. DNA extracted from 10 bacteria (*Vibrio harveyi, V. parahaemolyticus, V. vulnificus, V. tubiashi, V. alginolyticus, V. cholera, Streptococcus iniae, Tenacibaculum litopenaei, Pleisiomonas shigelloides*, and *Nocardia seriolae*) and DNA extracted from fish infected with 2 viruses (nervous necrosis virus (NNV) and infectious spleen and kidney necrosis virus (ISKNV)) were used as template. Details of the pathogens used in the specificity assay were described previously (Charoenwai et al., 2019). DNA extracted from clinically healthy Asian sea bass were also included in the assays. Positive and negative controls were reactions containing SDDV-infected fish DNA as template and water instead of DNA, respectively.

### 2.8 Detection of SDDV in clinical samples and analysis of viral loads

The qPCR condition was later employed for SDDV detection in DNA samples extracted from Asian sea bass tissues. 86 fish samples mentioned above were subjected to the detection. For quantitative analysis of SDDV viral loads in infected fish tissues, 3 fish samples collected in the year 2019 whose 8 different tissues had been preserved were used for investigation. Total DNA was extracted from eye, brain, fin, gills, kidney, liver, spleen, and muscle using a conventional phenol/ chloroform extraction and ethanol precipitation method. Viral copy numbers were calculated by extrapolating the Ct values to the standard curve generated as described above.

## 3. Results

### 3.1 SDDV qPCR condition optimization

Initial gradient PCR assays indicated that annealing temperature (Ta) ranging from 55 °C to 65 °C could be used for the designed qPCR primers. It was evidenced by the fact that SDDV infected fish yielded a specific band of 135 bp with similar band intensity (Fig. S2). However, when Ta of 60 and 63 °C was separately applied in SDDV qPCR assays, the melt curve analysis results revealed non-specific products probably being derived from primer-dimer formation (Fig. S2). Then, Ta of 63 °C was used but the primer concentration was reduced from 200 to 150 nM. Consequently, specific amplification was then obtained as demonstrated by a single uniform melting peak (Fig. S2). In conclusion, the optimized SDDV qPCR reaction of 20 μl contained 1x KAPA SYBR FAST master mix, 150 nM of each qSDDV-AF and qSDDV-AR primers, and 2 μl of DNA template. The cycling conditions run on ABI 7500 instrument consisted of denaturation at 95°C for 3 min followed by 40 cycles of 95°C for 3 s and 63°C for 30 s with subsequent melt curve analysis.

### 3.2 Amplification efficiency and detection sensitivity of SDDV qPCR assay

For detection sensitivity assay, an optimized SDDV qPCR condition above was run using 10-fold serially diluted SDDV control plasmid plus spiked fish DNA. It was found that the detection limit of the newly developed SDDV qPCR detection was 1 copy/μl template or 2 copies/reaction (Fig. 1a). The conditions were optimized after observing the uniform melting peaks (Tm 80.8 ± 0.15 °C) without detectable non-specific products (Fig. 1b). The resulting constructed standard curve (Fig. 1c) was then used for analysis of the amplification efficiency (E). With a slope of −3.115 (R^2^ 0.996), E of the SDDV qPCR developed in this study was 2.094, indicating a practical amplification assay.

**Figure 1.**
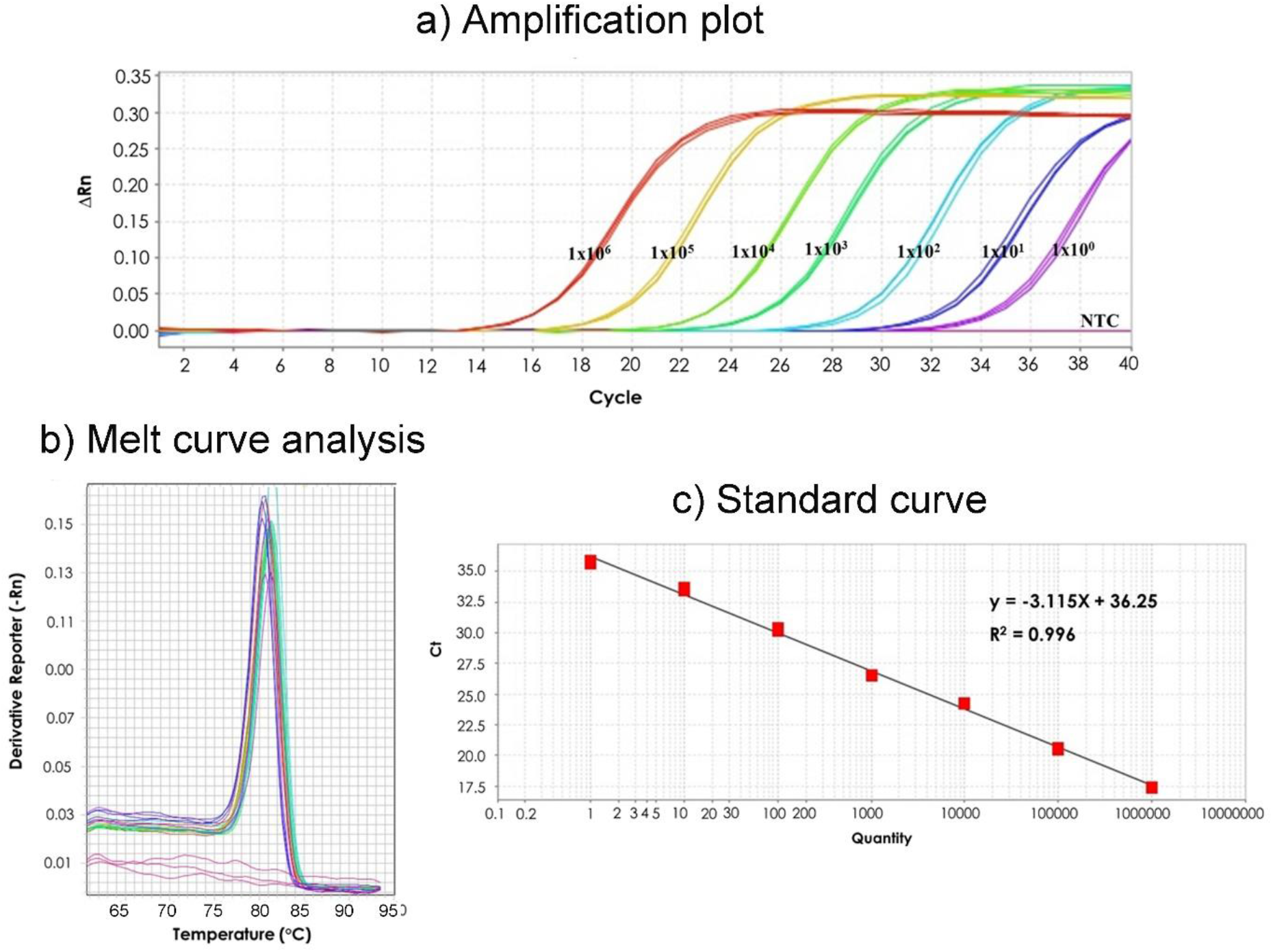
Development of SDDV qPCR assay. (a) Amplification plot of 10-fold serial dilution (1 to 10^6^ copies/μl) of positive control plasmid containing SDDV *ATPase* gene fragment. NTC, no template control. Experiment was performed in 3 replicates. (b) Melt curve analysis of the amplified amplicons obtained in (a) revealing uniform melt peaks of 80.8 ± 0.15 °C. (c) Standard curve derived by plotting between Ct values versus log concentration of plasmid copy number used in (a).

### 3.3 Reproducibility of SDDV qPCR assay

The reproducibility of the SDDV qPCR assay was characterized by analysis of intra-and interassay variations. The assays were performed using 10-fold serial dilutions of control plasmid containing SDDV *ATPase* gene fragment with 3 replicates. The results shown in Table 2 indicated that percent coefficient of variation (%CV) of the intra-and inter-assays were 0.24 to 0.70% with SD values ranging from 0.06-0.21 and 0.63 to 2.13% with SD values of 0.17-0.78, respectively. The analysis indicated the precision of results between different assays. It was also noted that the mean Ct value ranges were 35.76 to 17.38 (intra-assay) and 36.73 to 17.19 (inter-assay) when assayed with serial dilutions of control plasmid from 2×10^6^ to 2 plasmid copy numbers (Table 2).

**Table 2.** Reproducibility test of SDDV qPCR detection by intra-and inter-assay with 3 replicates each. 10-fold serial dilution of SDDV *ATPase* control plasmid was used as template

| Plasmid copy<br>number /reaction | Intra-assay |  |  | Inter-assay |  |  |
| --- | --- | --- | --- | --- | --- | --- |
|  | Mean Ct | SD | %CV | Mean Ct | SD | %CV |
| $2 \times 10^6$ | 17.38 | 0.08 | 0.45 | 17.19 | 0.23 | 1.36 |
| $2 \times 10^5$ | 20.57 | 0.12 | 0.56 | 20.51 | 0.18 | 0.91 |
| $2 \times 10^4$ | 24.25 | 0.06 | 0.24 | 24.11 | 0.21 | 0.85 |
| $2 \times 10^3$ | 26.52 | 0.10 | 0.38 | 26.35 | 0.17 | 0.65 |
| $2 \times 10^2$ | 30.23 | 0.21 | 0.70 | 30.12 | 0.29 | 0.97 |
| 20 | 33.62 | 0.19 | 0.56 | 33.84 | 0.21 | 0.63 |
| 2 | 35.76 | 0.17 | 0.48 | 36.73 | 0.78 | 2.13 |

### 3.4 Specificity test of SDDV qPCR detection method

DNA samples prepared from 12 pathogens commonly found in aquatic animals consisting of 10 bacteria and 2 viruses were subjected to specificity test. DNA extracted from SDDV-infected fish and a clinically healthy fish was used in positive and negative control reactions, respectively. The result shown in Fig. 2 indicated that specific detection was obtained from only the SDDV-infected sample and no cross amplification with other pathogens or DNA from healthy fish was observed.

**Figure 2.**
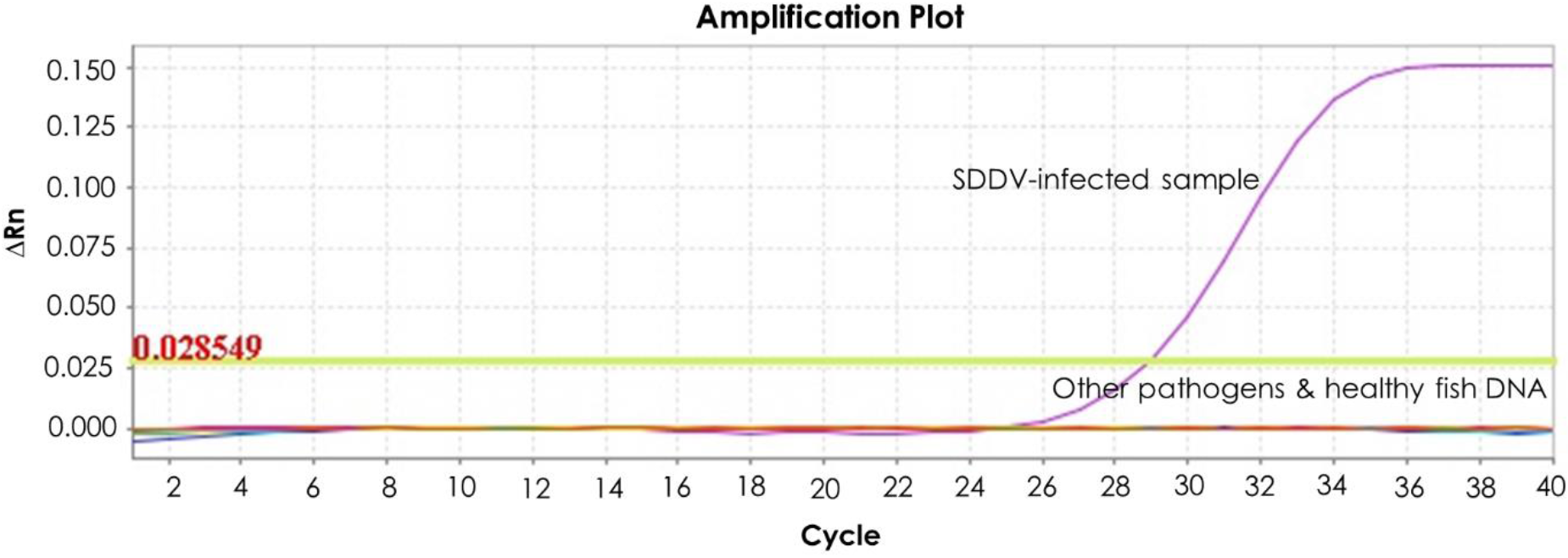
Specificity test of the SDDV qPCR assay. The test was performed using DNA extracted from an SDDV-infected fish, a healthy fish, and 12 bacterial and viral pathogens commonly found in aquatic animals. Details of pathogens are in materials and methods section.

### 3.5 Detection of SDDV in clinical samples

The established SDDV qPCR condition was employed to investigate the presence of SDDV in the samples collected from different farms in 2016-2019 (Table 1). A total number of 86 fish samples were obtained from 34 scale drop diseased or unknown diseased fish and 52 healthy looking Asian sea bass. This set of samples was previously tested using semi-nested PCR method (Charoenwai et al., 2019) and found that all the diseased fish were SDDV positive while the clinically healthy fish were tested negative for SDDV. Interestingly, the qPCR detection result shown in Table 1 indicated that 30 out of the 52 healthy looking fish were SDDV positive (Ct values ranging from 29.95-37.02). As expected, all of the 34 sick fish were confirmed to be SDDV infected by qPCR (Ct values ranging from 19.07-28.73). Five representative 135-bp amplicons from positive samples obtained from 5 farms (1, 8, 9, 10, and 11) were subjected to cloning and sequencing. DNA sequence analysis revealed high identity (98.52-100%) to the sequence of SDDV Singapore isolate deposited in the database (KR139659)(Fig. 3).

**Figure 3.**
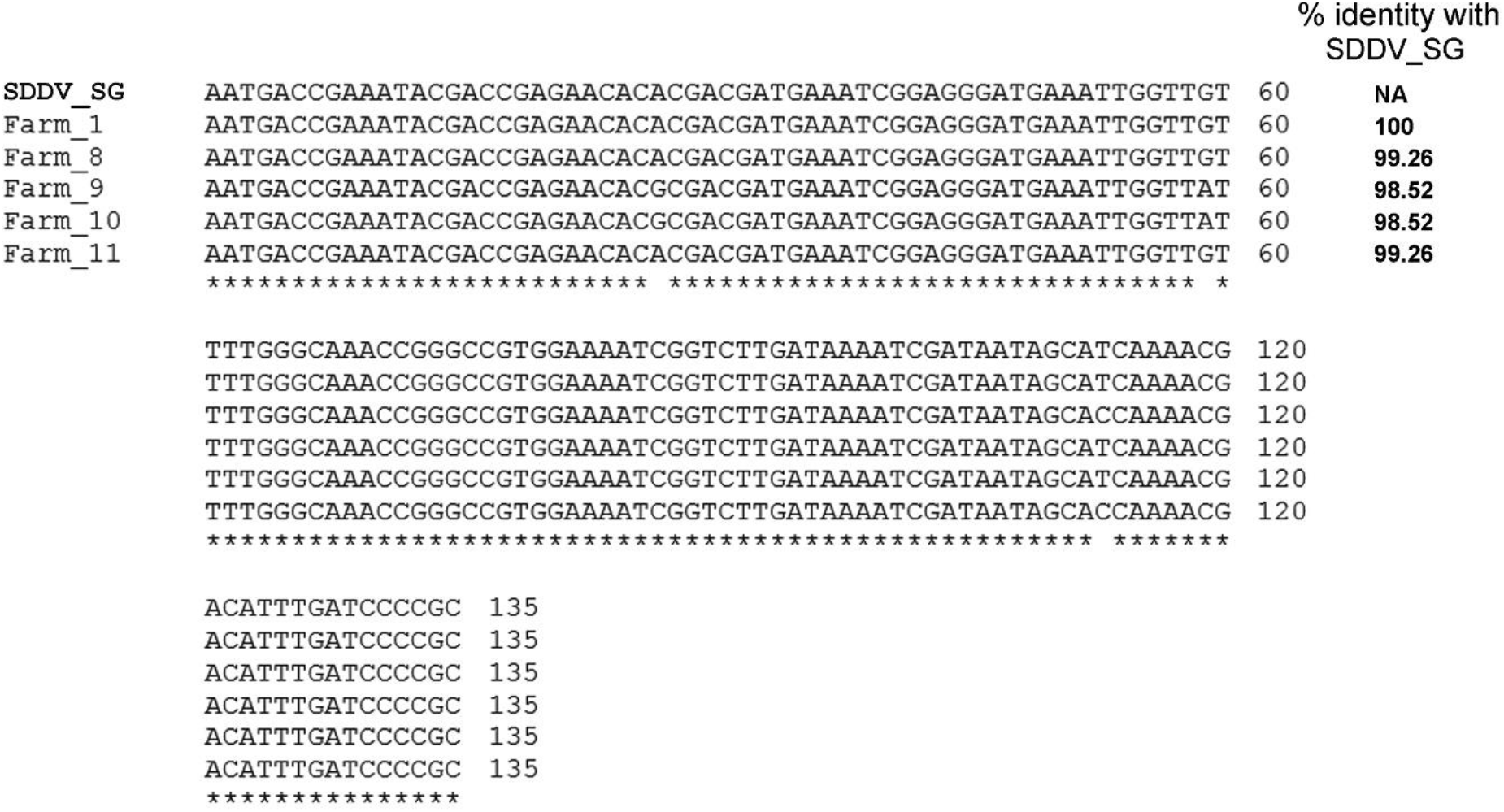
Nucleotide sequence alignment of 135-bp SDDV *ATPase* partial gene fragments. Sequences from representative amplicons from SDDV qPCR positive test samples obtained in this study (5 different farms) were compared to that of SDDV Singapore isolate retrieved from GenBank database (KR139659). % identity to the Singapore isolate is shown on the right panel.

### 3.6 Detection of SDDV in different fish tissues

In this study, 3 SDDV-infected fish collected in 2019 were subjected to investigation of tissue tropism of SDDV infection. DNA from 8 different tissues including eye, brain, fin, gills, kidney, liver, spleen, and muscle of individual fish were tested using our newly developed SDDV qPCR protocol. The results showed that all of the tested tissues were infected with SDDV with the viral loads ranging from 8.0×10^2^ to 6.8×10^4^ viral copies/200 ng of of DNA template (Table 3). Brain, fin, gills, and muscle seem to have relatively higher viral load (6.4×10^3^ to 6.8×10^4^) when compared to the rest of other organs (8.0×10^2^ to 3.7×10^4^).

**Table 3.**
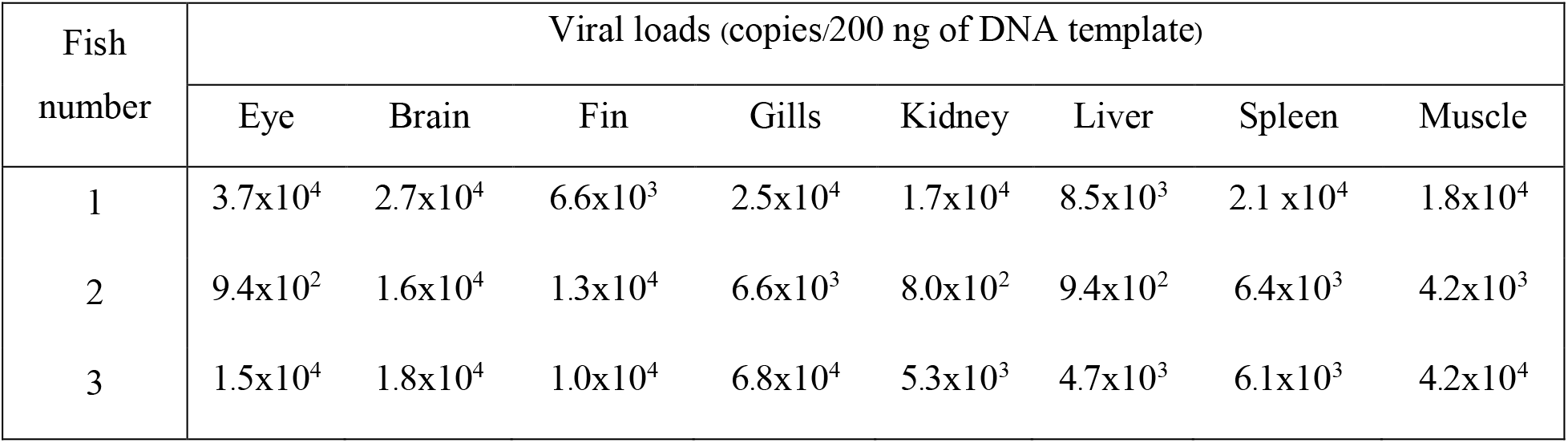
Analysis of SDDV viral loads in different fish tissues

| Fish<br>number | Viral loads (copies/200 ng of DNA template) |  |  |  |  |  |  |  |
| --- | --- | --- | --- | --- | --- | --- | --- | --- |
|  | Eye | Brain | Fin | Gills | Kidney | Liver | Spleen | Muscle |
| 1 | $3.7 \times 10^4$ | $2.7 \times 10^4$ | $6.6 \times 10^3$ | $2.5 \times 10^4$ | $1.7 \times 10^4$ | $8.5 \times 10^3$ | $2.1 \times 10^4$ | $1.8 \times 10^4$ |
| 2 | $9.4 \times 10^2$ | $1.6 \times 10^4$ | $1.3 \times 10^4$ | $6.6 \times 10^3$ | $8.0 \times 10^2$ | $9.4 \times 10^2$ | $6.4 \times 10^3$ | $4.2 \times 10^3$ |
| 3 | $1.5 \times 10^4$ | $1.8 \times 10^4$ | $1.0 \times 10^4$ | $6.8 \times 10^4$ | $5.3 \times 10^3$ | $4.7 \times 10^3$ | $6.1 \times 10^3$ | $4.2 \times 10^4$ |

## 4. Discussion

Molecular diagnosis of SDDV from clinically sick fish could employ only a single PCR assay due to high viral load in the infected tissues (Senapin et al., 2019). However, single PCR or even nested PCR assays sometimes resulted in false negative detection for the samples that have viral load under detection limit of the methods. In aquatic animal culture, there is possibly a large proportion of viral infections in subclinical form that did not exhibit abnormal clinical signs (Senapin et al., 2018; Jeamkunakorn et al., 2019). Thus, accurate and sensitive diagnosis for these samples requires the assays which can detect as low as 1 −2 viral copies per reaction to avoid false negative results, especially for live fish which are being translocated and destined for aquaculture.

The newly established qPCR assay described in this study is highly specific to SDDV and could detect down to 2 copies per reaction which is 100 times more sensitive than our previous sem-inested PCR and LAMP protocols (Charoenwai et al., 2019; Dangtip et al., 2019), and 25 times more sensitive than a previously probe-based qPCR detection (de Groof et al., 2015). When applied to detection of clinically healthy fish from unaffected farms, it was revealed that ~57.7g% (30/52) samples previously tested negative by semi-nested PCR were positive by qPCR, indicating that the newly established qPCR is suitable for detection of SDDV from subclinically infected fish populations. Thus, we recommend the SDDV *ATPase* qPCR method should be considered for active surveillance program and quarantine inspection to prevent widespread of the pathogen. For farmers, this sensitive qPCR method might be useful for selection of SDDV-tested negative seeds before stocking to avoid the risk of disease outbreak.

Analysis of viral load of SDDV in different tissues of the infected fish revealed presence of the virus in all 8 tested tissues (eye, brain, fin, gills, kidney, liver, spleen, and muscle). This suggested that SDDV caused systemic infection and the viral load seemed to be relatively higher in the brain, fin, gills, and muscle when compared to other organs. This finding might suggest potential use of fin and gills as target organs for non-lethal detection of SDDV. However, it is not certain whether these organs are suitable for detection in case of subclinically infected fish with presumably low viral load. Thus, further investigation on comparative viral load of SDDV in different infection levels is required as basic knowledge for development of non-lethal sampling methods, especially for high value broodfish.

Moreover, it is worthwhile to investigate further whether SDDV invades the reproductive organs of Asian sea bass broodfish and could vertically transmit to the offspring as occurred in the case of tilapia lake virus (TiLV), a newly emerging virus of tilapia (Dong et al., 2020; Yamkasem et al., 2019). Understanding the route of disease transmission in combination with highly sensitive detection method is vital for effective disease management and preventing potential widespread.

In conclusion, this study has developed a highly sensitive qPCR method for the specific detection of SDDV in Asian sea bass with detection limit of 2 viral copies per reaction. The method was able to detect SDDV in both clinically sick fish and inapparently (asymptomatic) infected fish. Thus, qPCR protocol developed in this study might be a useful tool for SDDV diagnosis and surveillance.

## Acknowledgements

This work was supported by BIOTEC fellow’s research grant P-19-50170. S. Sriisan is the recipient of a Thailand Graduate Institute of Science and Technology (TGIST), NSTDA scholarship. The authors would like to thank Ms. O. Charoenwai and Mr. W. Meemetta for skilled technical assistance.

---

**Figure S1.**
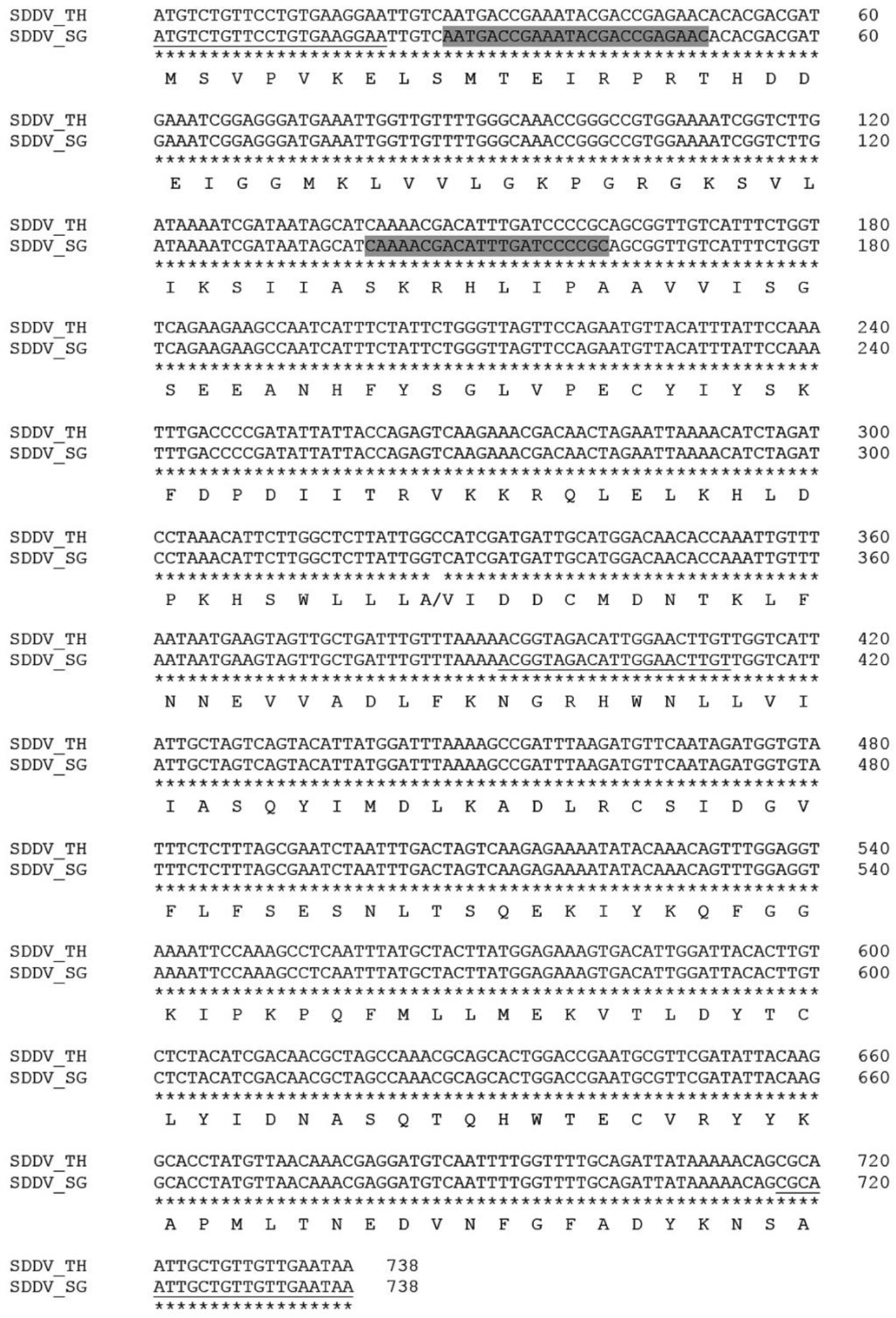
Position of SDDV detection primers. Open reading frame sequences of SDDV *ATPase* gene from Thai (TH, MH152407) and Singapore (SG, KR139659) isolates retrieved from GenBank database were compared. Primers used in qPCR assay developed in this study and in semi-nested PCR described in Charoenwai et al. (2019) were gray-highlighted and underlined, respectively.

**Figure S2.**
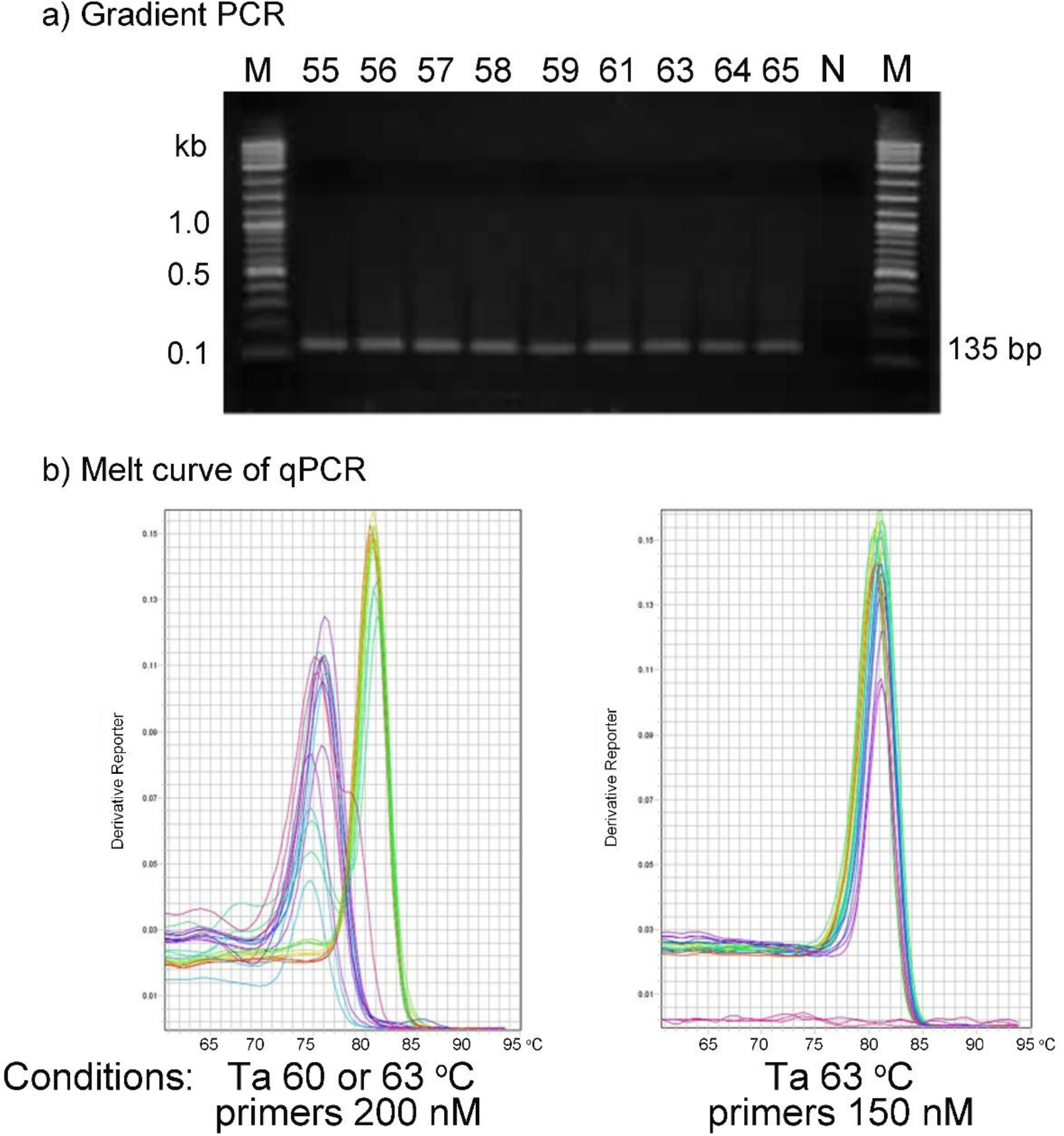
Optimizing the SDDV qPCR conditions. (a) Gradient conventional PCR using Ta at 55-65 °C were performed with reactions containing SDDV-infected fish DNA and qSDDV-AF and qSDDV-AR primers. N, no template control. (b) Melt curve analysis of qPCR assays using Ta at 60 or 63 °C and primer concentration of 200 nM showing non-specific products (left) and conditions using Ta at 63 °C and primer concentration of 150 nM revealing uniform melt peaks (right).

